# VMDance: An Efficient and Interactive Molecular Dynamics Clustering Plugin for VMD

**DOI:** 10.64898/2026.11.24.769027

**Authors:** Diego E. B. Gomes, Jherome Brylle Woody A. Santos, Krisztina A. Zsigmond, Vicky Jung, Jerwin C. Quirante, Rafael C. Bernardi, Ramón Alain Miranda-Quintana

**Affiliations:** Department of Physics, Auburn University, Auburn, AL, USA; Department of Chemistry and Quantum Theory Project, University of Florida, Gainesville, FL, USA

**Keywords:** molecular dynamics, trajectory clustering, *n*-ary similarity, conformational analysis, scalable analysis, molecular visualization, VMD

## Abstract

Clustering is the standard route from a molecular dynamics trajectory to a small set of interpretable conformational states, yet it is usually performed outside the molecular viewer, leaving cluster assignments to be exported, tracked, and mapped back onto the structures by hand. We present VMDance, an open-source plugin that brings the MDANCE family of *n*-ary similarity clustering methods into VMD. Through a graphical interface backed by a C++ implementation, it provides *k*-means NANI, HELM, eQual, and DIVINE clustering, together with PRIME representative-structure selection. These methods scale linearly with the number of frames and support deterministic initialization, so partitions can be reproduced exactly and recomputed interactively. Cluster assignments, quality metrics, and representative structures are linked to the loaded trajectory, keeping the analysis of conformational states alongside the structures themselves. VMDance is freely available at https://github.com/diegoenry/vmd-mdance-plugin.

## 1 Introduction

Proteins change conformation as they bind partners, catalyze reactions, and transport solutes. Molecular dynamics (MD) simulations can resolve the motions underlying these processes. ^1,2^ In HIV-1 protease, simulations captured the opening and closing of two flaps that control access to the active site. ^3^ In the glycerol-3-phosphate antiporter GlpT, a larger rearrangement exposes the same substrate-binding site first to one side of the membrane and then to the other. ^4^

At protein–protein interfaces, the relevant structural differences often concern how the partners bind or how they separate. Dynamic network analysis distinguished two proposed binding orientations for an Affibody on PD-L1, ^5^ and the conformation of Hsp60 influenced where its partner MIX was predicted to dock. ^6^ Binding geometry also affects the response to force. Two cohesin–dockerin binding modes ruptured at different forces, ^7^ while the adhesin SdrG followed distinct dissociation pathways at low and high forces. ^8^ Across SARS-CoV-2 variants, equilibrium affinity for ACE2 did not consistently predict the mechanical stability of the interface. ^9^

Clustering helps identify recurring conformations by grouping structurally similar frames, quantifying their populations in a trajectory, and selecting representative structures for inspection. ^10–12^ The resulting groups depend on the atom selection, the algorithm, and parameters such as the number of clusters (*k*).^10,13^ Inspecting representative structures alongside cluster quality indices in VMD lets researchers see whether a partition resolves the motion of interest or primarily reflects flexibility elsewhere, then refine the analysis accordingly.

Computational cost and reproducibility become important when clustering a trajectory repeatedly. Some hierarchical agglomerative clustering (HAC) implementations use a full matrix of pair-wise root-mean-square deviations (RMSDs). Constructing and storing this matrix requires time and memory that scale quadratically with the number of frames. ^10,14,15^ Conventional *k*-means avoids this matrix by comparing frames with cluster centroids. ^11^ With stochastic initialization methods such as *k*-means++, different random seeds can yield different cluster assignments and populations. ^16^

*n*-ary similarity allows structural dispersion to be computed from coordinate-wise sums and squared sums over an ensemble rather than from a full pairwise distance matrix.^17–20^ For aligned Cartesian coordinates, the mean squared deviation (MSD) of a set of *N* frames can therefore be calculated in *O*(*N*) time for a fixed number of atoms, without constructing a pairwise distance matrix. The same quantities rank frames by structural centrality, again without storing pairwise distances.

The Molecular Dynamics Analysis with *N* -ary Clustering Ensembles (MDANCE) package builds on these quantities. ^21^ It includes *k*-means NANI, ^22,23^ which selects initial centroids for *k*-means deterministically; HELM, ^24^ which merges preclusters without constructing a full frame-by-frame distance matrix; eQual, ^25^ which uses a radial-threshold procedure; DIVINE, ^26^ which builds a divisive hierarchy; and PRIME,^27^ which selects a representative structure from a clustered ensemble.

Molecular visualization is essential for interpreting clustering results in the context of the underlying structures. Although clustering reduces a trajectory to a set of conformational states and representative structures, the analysis is often performed separately from the molecular viewer. Cluster assignments must therefore be mapped back to the trajectory, requiring additional data handling and careful tracking of frame indices. This separation becomes particularly inconvenient when repeated clustering runs are needed to evaluate different algorithms and parameters. Clustering is available in analysis packages such as CPPTRAJ,^28^ MDAnalysis, ^29^ MDTraj,^30^ and PyEMMA, ^31^ and in standalone tools such as TTClust. ^32^ Within VMD, the built-in `measure cluster` command ^33^ performs cutoff-based clustering that has been shown to follow the algorithm of Daura et al., ^34^ and the Clustering plugin ^35^ provides a graphical interface to this command and to results imported from external programs. None of these provides the *n*-ary methods inside the molecular viewer.

Here, we present VMDance, a plugin that integrates MDANCE analysis, evaluation, and selection of representative frames with the molecular visualization capabilities of VMD. ^33^ Using the C++ implementation CPP-MDANCE, VMDance connects cluster assignments and quality scores to the loaded trajectory. The interface emphasizes atom and frame selection, clustering parameters, and the inspection of representative structures, allowing users to evaluate results alongside the molecular structures. The plugin is described in Section 2, the available methods in Section 3, and an application to a trajectory in Section 4.

## 2 Software and Usage

### 2.1 Architecture

VMDance is written in Tcl, VMD’s scripting language, and is organized around a core that does not use the Tk graphical toolkit and is shared by two front ends (Figure 1): the graphical window, which is loaded only when it is opened, and the `vmdance` command, which provides the same functions to scripts and to VMD running in text mode (`vmd -dispdev text`). The core reads frames from the loaded molecule, prepares the coordinates, runs the backend, stores each clustering together with its quality measures, and colors the clusters in the VMD 3D window. The window holds no results of its own and is updated by the core whenever a run, scan, or selection changes. A single parameter table defines every algorithm option and generates the command options, the window fields, their validity checks, and the arguments passed to the backend. A clustering job does not change the current frame or coordinates of the loaded molecule, and a failed or canceled job leaves the previous result unchanged.

**Figure 1.**
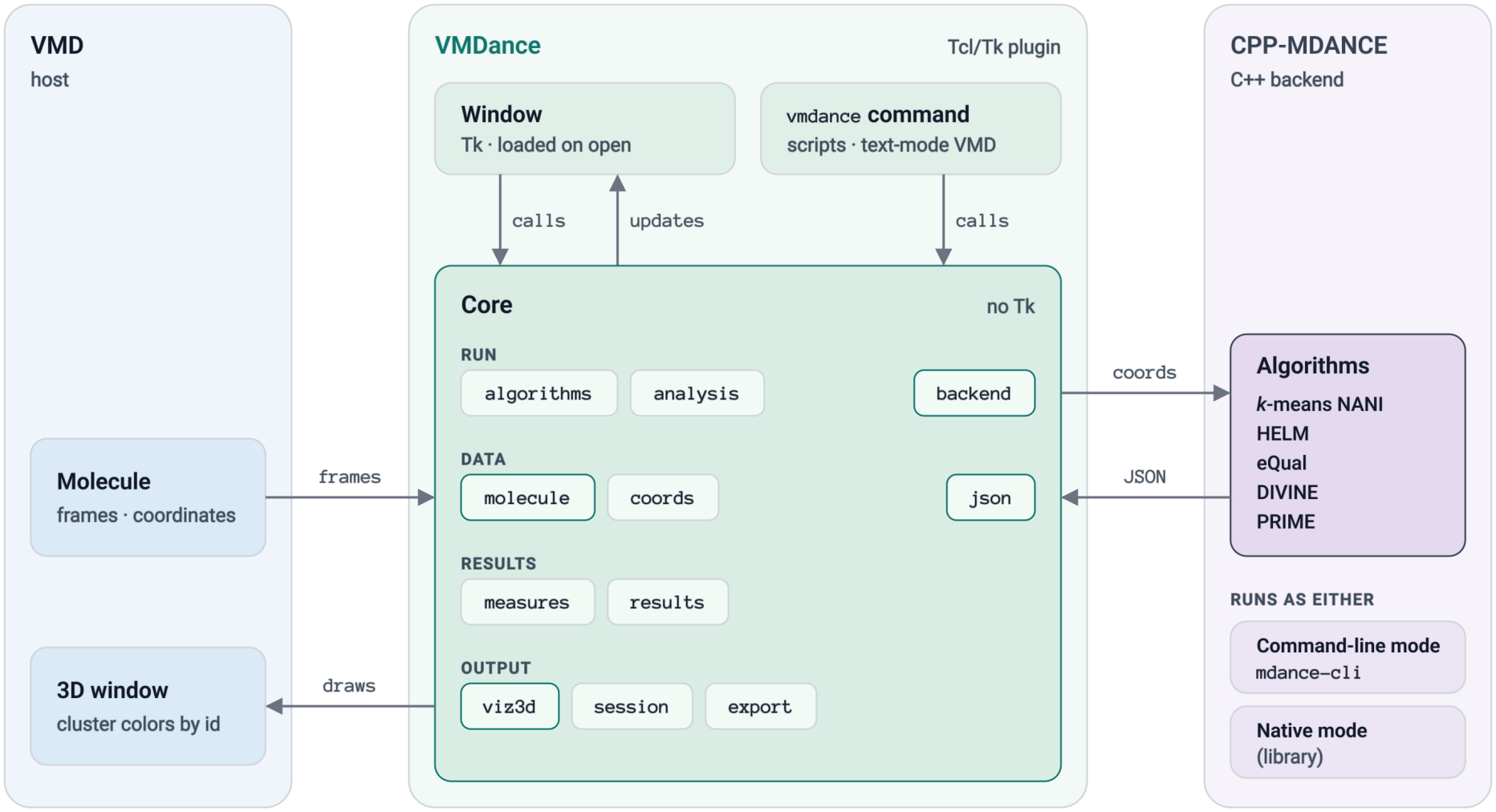
Architecture of VMDance. The core is shared by the graphical window and the `vmdance` command. It reads frames from the molecule loaded in VMD, passes the coordinates to CPP-MDANCE, reads the results, and colors the clusters in the 3D window. CPP-MDANCE runs either as the `mdance-cli` program (command-line mode) or as a library inside VMD (native mode).

The core communicates with the backend through two execution modes. In native mode, a Tcl extension passes coordinate data directly to the C++ library in memory, avoiding temporary files and subprocess creation. In command-line mode, the plugin writes the coordinates to a temporary CSV file, executes `mdance-cli`, and reads the resulting JSON output. The command-line mode supports progress reporting and cancellation, whereas native calls execute within the VMD process.

### 2.2 Interface

The window has three control regions (Figure 2A). The global toolbar (region 1; Figure 2B) provides shortcuts for opening and saving sessions, exporting results and figures, running or canceling clustering calculations, initiating scans, accessing plots, and controlling molecular visualization. Settings and help are also available from this toolbar. The primary navigation tabs (region 2; Figure 2C) consist of Setup, Results, Scans, Tools, and 3D. Setup provides molecule and atom selection, frame-range controls, and algorithm parameters, with atom and frame counts updated as selections change. The Results tab reports cluster assignments, populations, MSD values, quality scores, and representative frames, and retains runs for comparison. Scans provides elbow scans and parameter sweeps; Tools contains PRIME, extended-similarity analysis, and frame-selection functions; and 3D provides structural visualization controls.

**Figure 2.**
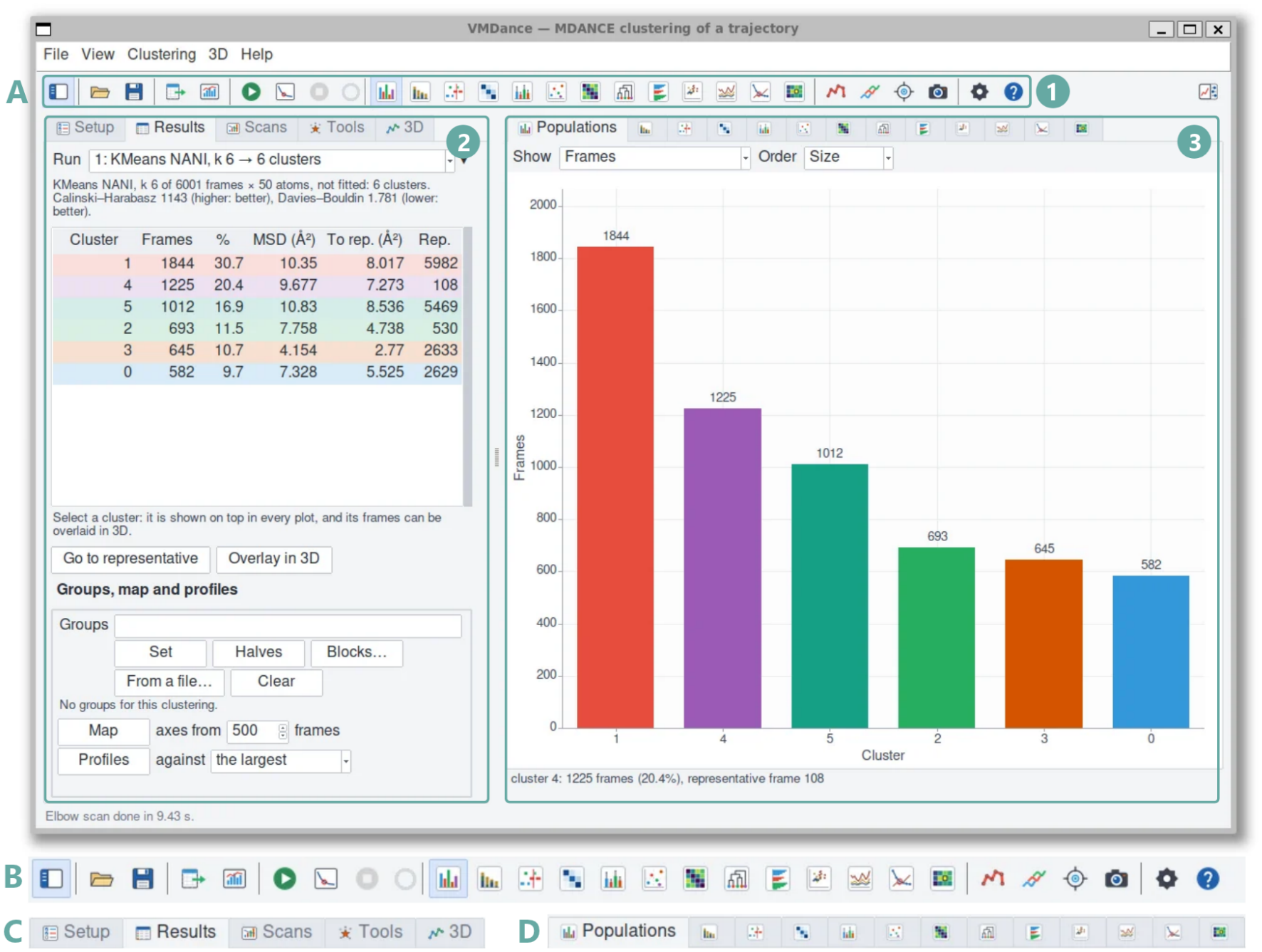
VMDance graphical user interface after a *k*-means NANI run (`comp_sim` seeding, *k* = 6) on the tau peptide. (A) Full interface showing the global toolbar (region 1), primary navigation tabs (region 2), and plot navigation toolbar (region 3). (B) Enlarged view of the global toolbar. (C) Enlarged view of the primary navigation tabs. (D) Enlarged view of the plot navigation toolbar.

The plot navigation toolbar (region 3; Figure 2D) gives access to 13 diagnostic views: Populations, Make-up, Timeline, Transitions, Residence, Compactness, Distances, Tree, Silhouette, Map, Profiles, Elbow, and Sweep. These display cluster populations, assignments over time, lag-dependent transitions, residence times, structural dispersion and separation, relationships between clusters, and clustering quality across parameter choices. Cluster assignments are also mapped to VMD’s User field for coloring, allowing cluster selections and representative structures to be examined in the molecular view using the same colors as the plots.

### 2.3 Coordinate Handling and Frame Mapping

The plugin extracts Cartesian coordinates from the selected atoms and frames for processing by CPP-MDANCE. Because the clustering methods operate on Cartesian coordinates, structural alignment is necessary to remove rigid-body translation and rotation. ^36^ Trajectories can be aligned beforehand using VMD’s RMSD Trajectory Tool, or optionally fitted during coordinate extraction using a specified atom selection. The latter transforms only the coordinates passed to the backend, leaving the loaded trajectory unchanged.

VMDance maintains an explicit mapping between the analyzed frames and their original VMD frame numbers. This mapping is preserved throughout clustering, visualization, and export, even when only a subset of frames is analyzed. Frames outside the selected range remain unassigned, preventing their accidental inclusion in cluster representations.

### 2.4 Exports

A completed run can be written out as per-frame labels in CSV, one PDB or DCD per cluster medoid, one trajectory file per cluster, the top-*N* frame indices of each cluster, or a session file containing the result with its parameters and frame map for later reopening. Any plot exports its underlying data as CSV and the figure itself as PostScript or PNG, composed at a fixed figure size independent of the window size.

## 3 Methods

All methods in VMDance operate on the Cartesian coordinates of a user-defined atom selection. Each frame is represented by a vector of 3*M* coordinates, where *M* is the number of atoms. Each structure may be aligned to a reference frame before clustering, as described in Section 2.3. The methods rely on two *n*-ary quantities: the MSD of a set of frames, which measures how compact the set is, and the complementary mean squared deviation (cMSD) of each frame, which ranks frames based on their centrality. Figure 3 outlines the workflow of each method, and Table 1 lists their main parameters. Derivations and validation are shown in the original publications.

**Table 1:** Clustering and analysis methods available in VMDance.

| **Method** | **Type** | **Principal user parameters** | **Refs.** |
|---|---|---|---|
| *k*-means NANI | Partitional | *k*; seeding; seed fraction | 22, 23 |
| HELM | Agglomerative hybrid | *k*; initial clusters; merge criterion | 24 |
| eQual | Radial threshold | MSD threshold; seeding; minimum size | 25 |
| DIVINE | Divisive | *k*; split criterion; seeding | 26 |
| PRIME | Post-processing | Similarity index; trimming; weighting | 27 |

**Figure 3.**
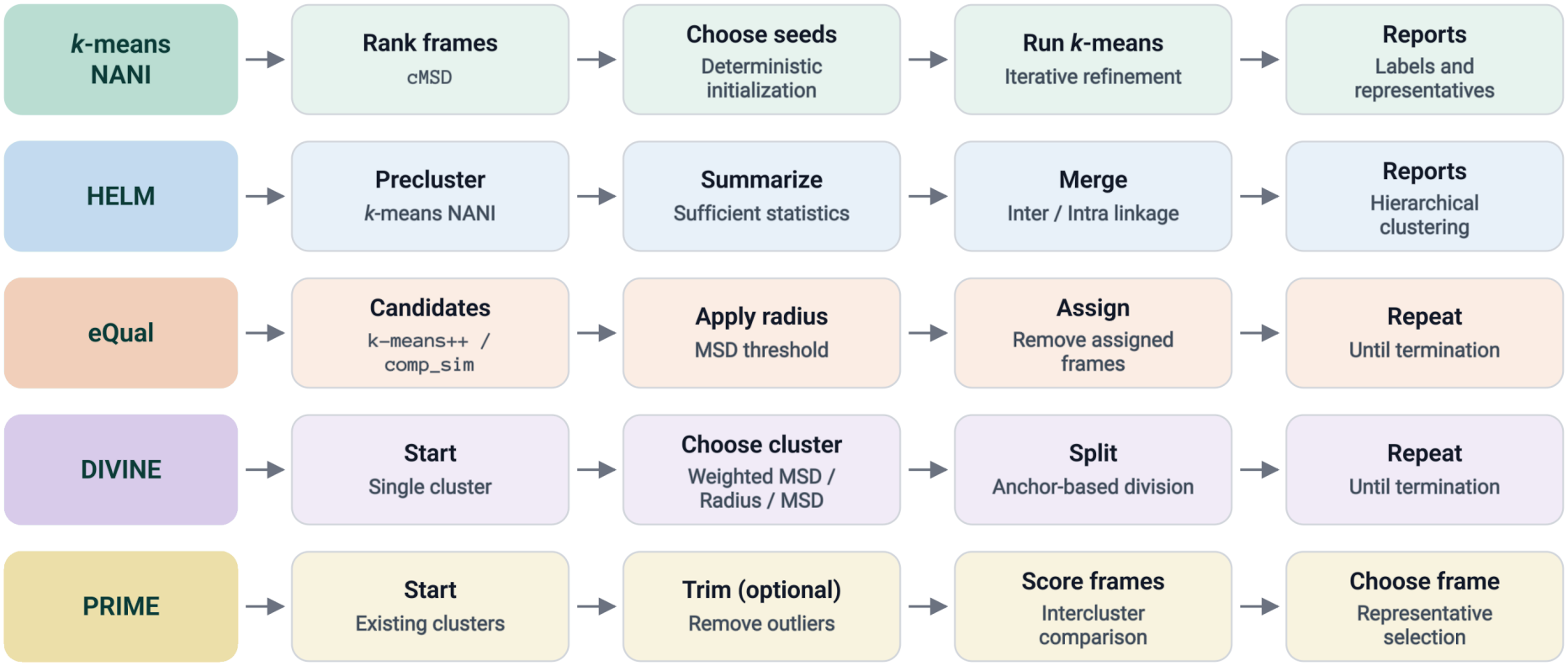
Workflows of the methods available in VMDance.

### 3.1 *k*-means NANI

*k*-means *N* -ary Natural Initiation (NANI) ^22^ addresses the sensitivity of *k*-means to its starting seeds. Instead of drawing the initial centroids at random, NANI ranks all frames by cMSD and selects seeds that lie in densely populated regions of the conformational space but are diverse. The assignment and update steps of *k*-means are then carried out as usual. Under deterministic seeding and fixed input conditions, repeated runs will always produce identical cluster assignments and populations.

In the original implementation (`comp_sim`), seeds were chosen one at a time from the top fraction of the cMSD ranking, each time taking the frame least similar to the seeds already selected. Two stratified variants were later introduced as faster alternatives to this iterative search. ^23^ The first, `strat_all`, divides the ranked list of frames into equal strata and takes the highest-ranked frame of each; the second, `strat_reduced`, does the same within the densest fraction of frames only.

### 3.2 HELM

The Hierarchical Extended Linkage Method (HELM) ^24^ combines the efficiency of *k*-means with the flexibility of agglomerative clustering. The trajectory is first divided with *k*-means NANI into small microclusters, with a number of initial clusters greater than the desired number of clusters (e.g., 30 to 60). Each microcluster is then summarized by three quantities: its number of frames, the coordinate-wise sum of its frames, and the coordinate-wise sum of their squares. These summaries give the MSD of any union of two clusters directly, so a matrix of dissimilarities between the microclusters can be built and the clusters merged step by step without returning to individual frames.

Two *n*-ary merging criteria are available: `intra`, the MSD of the union between two clusters, and `inter`, the average MSD between their members. The method also allows the trimming of high-MSD or sparsely populated clusters, which could add noise, before merging. The result is a merge hierarchy that can be cut at any *k*.

### 3.3 eQual

Extended Quality (eQual) ^25^ clustering is a linear-scaling version of the radial threshold clustering introduced by Daura and co-workers. ^37^ In the original algorithm, every frame proposes a cluster made of all frames within an RMSD cutoff, ^34^ which requires the full pairwise matrix. eQual instead draws a few candidate seeds at each iteration, either with *k*-means++ or by complementary similarity, and lets each candidate collect the frames that lie within a user-defined MSD threshold. The most populated candidate becomes a cluster, its frames are removed, and the procedure repeats until no frames remain.

When two candidates contain the same number of frames, the one with the lower MSD is kept, so the result does not depend on the order of frames in the trajectory. With complementary similarity seeding, the procedure is fully deterministic. With *k*-means++ seeding, the results depend on the random seed. The number of clusters, *k*, is not fixed in advance but follows from the threshold, which could be convenient when states are naturally defined by a maximum structural deviation. Optional filters discard clusters that exceed a similarity threshold or are too small.

### 3.4 DIVINE

DIVIsive *N* -ary Ensembles (DIVINE) ^26^ builds a clustering hierarchy from the top down. All frames start within a single cluster, and at each step one cluster is chosen and divided into two. By default, DIVINE splits the cluster with the largest weighted MSD, the product of MSD and population, which favors large heterogeneous clusters over small groups of outliers. The unweighted MSD and the cluster radius are also available as selection criteria. The two seeds of each split are obtained with NANI (*k* = 2) or from outlier-based anchors, which are followed by a short *k*-means refinement. Clusters are split continuously until the requested *k* is reached or every frame forms its own cluster. Quality scores, populations, and MSD values are recorded after each split, so a single run recovers all resolutions while preserving the hierarchical relationships between clusters. Unlike conventional *k*-means, which produces a flat partition at a specified *k*, DIVINE constructs a hierarchy that tracks how clusters progressively split into distinct conformational states.

### 3.5 PRIME

Protein Retrieval via Integrative Molecular Ensembles (PRIME) ^27^ is not a clustering method but a post-processing step that can follow any clustering protocol. Conventional representative selection often relies on the centroid or medoid of the most populated cluster, without considering information from the remaining clusters. PRIME instead scores each frame from the most populated cluster by its similarity to the remaining clusters, weighting each cluster by its population, after optionally removing the most outlying frames.

The comparison can be made with every frame of the other clusters, with each cluster as a whole through extended similarity, or with the medoid or outlier of each cluster.

### 3.6 Cluster Quality

For every partition, VMDance reports the Calinski–Harabasz index (CHI), ^38^ also known as the pseudo-*F* statistic, which compares between-cluster and within-cluster dispersion, and the Davies– Bouldin index (DBI), ^39^ which compares within-cluster spread with between-cluster separation. Higher CHI and lower DBI values generally indicate better separation. These indices may favor different choices of *k* and should be considered alongside cluster populations and spread. The plugin also reports silhouette values ^40^ as a frame-level assessment. Scanning these indices over the number of clusters helps users to choose the optimal *k*, either from the global extremum or from extrema of their second derivatives, which mark partitions that are more stable than their neighbors.

## 4 Case Study: Clustering a Tau Peptide Trajectory

We illustrate the workflow on a 6,001-frame trajectory of the tau peptide from a replica-exchange MD simulation, ^41^ using publicly available topology and trajectory files. ^34,42^ Clustering used the backbone N, C*α*, C, O, and H atoms of residues Lys2–Asp11 (50 atoms; `resid 3 to 12` in the topology file), following Daura and Conchillo-Solé. ^43^ The trajectory was aligned to the 1000^th^ frame before clustering. All timings were measured with the CPP-MDANCE command-line backend (`mdance-cli`) on a single core of an AMD Ryzen 7 8845HS.

### 4.1 Choosing *k*

The trajectory was clustered with *k*-means NANI using `comp_sim` seeding with a 10% seed fraction, and an elbow scan over *k* = 3–10 (Table 2) was used to select the number of clusters. Partitions with *k* = 2 were not considered: although DBI is lowest there (1.5371), a two-state split is trivial for an ensemble known to sample more than two conformational states. ^22^ CHI decreases monotonically with *k*, and DBI reaches its minimum at *k* = 6 (1.7809), which also has the second-largest DBI second derivative, after *k* = 8 (Figure S1). This partition was used in the following analysis; extending the scan to *k* = 30 does not change the minimum for *k ≥* 3 (Table S1).

**Table 2:** Elbow scan of *k*-means NANI (comp_sim seeding, 10% seed fraction) on the tau peptide (6,001 frames). CHI, Calinski–Harabasz index (higher is better); DBI, Davies–Bouldin index (lower is better); Avg MSD, unweighted mean of per-cluster mean squared deviation. The selected partition is in bold.

| *k* | CHI | DBI | Avg MSD | Cluster sizes |
|---|---|---|---|---|
| 3 | 1750.28 | 1.8168 | 10.881 | 2338, 2337, 1326 |
| 4 | 1434.61 | 1.9856 | 10.122 | 1765, 1690, 1343, 1203 |
| 5 | 1298.01 | 1.7952 | 9.026 | 1809, 1354, 1191, 833, 814 |
| **6** | **1142.79** | **1.7809** | **8.349** | **1844, 1225, 1012, 693, 645, 582** |
| 7 | 1051.14 | 1.9452 | 8.107 | 1167, 1130, 1052, 918, 638, 614, 482 |
| 8 | 967.30 | 1.9277 | 7.820 | 1097, 999, 812, 800, 755, 645, 478, 415 |
| 9 | 882.01 | 2.0898 | 7.920 | 804, 724, 723, 719, 690, 665, 662, 613, 401 |
| 10 | 842.80 | 1.9944 | 7.649 | 829, 698, 691, 674, 609, 545, 536, 498, 464, 457 |

### 4.2 Reading a Partition

At *k* = 6, the clusters hold between 9.7% and 30.7% of the frames, with per-cluster MSDs between 4.15 and 10.83 Å^2^ (CHI 1142.79, DBI 1.7809). For every partition in Table 2, the CHI, DBI, average MSD, cluster populations, and per-cluster MSD values agree with those from the Python implementation of MDANCE. The run completes in 0.40–0.44 s (wall time over five runs of `mdance-cli`, including reading the input and writing the results). Cluster 3 (645 frames, 10.7%; clusters are numbered from 0) has an MSD of 4.15, less than half that of the three most populated clusters (9.68–10.83), which marks it as the most compact state in the ensemble. The representative structures of each cluster are shown in Figure 4.

**Figure 4.**
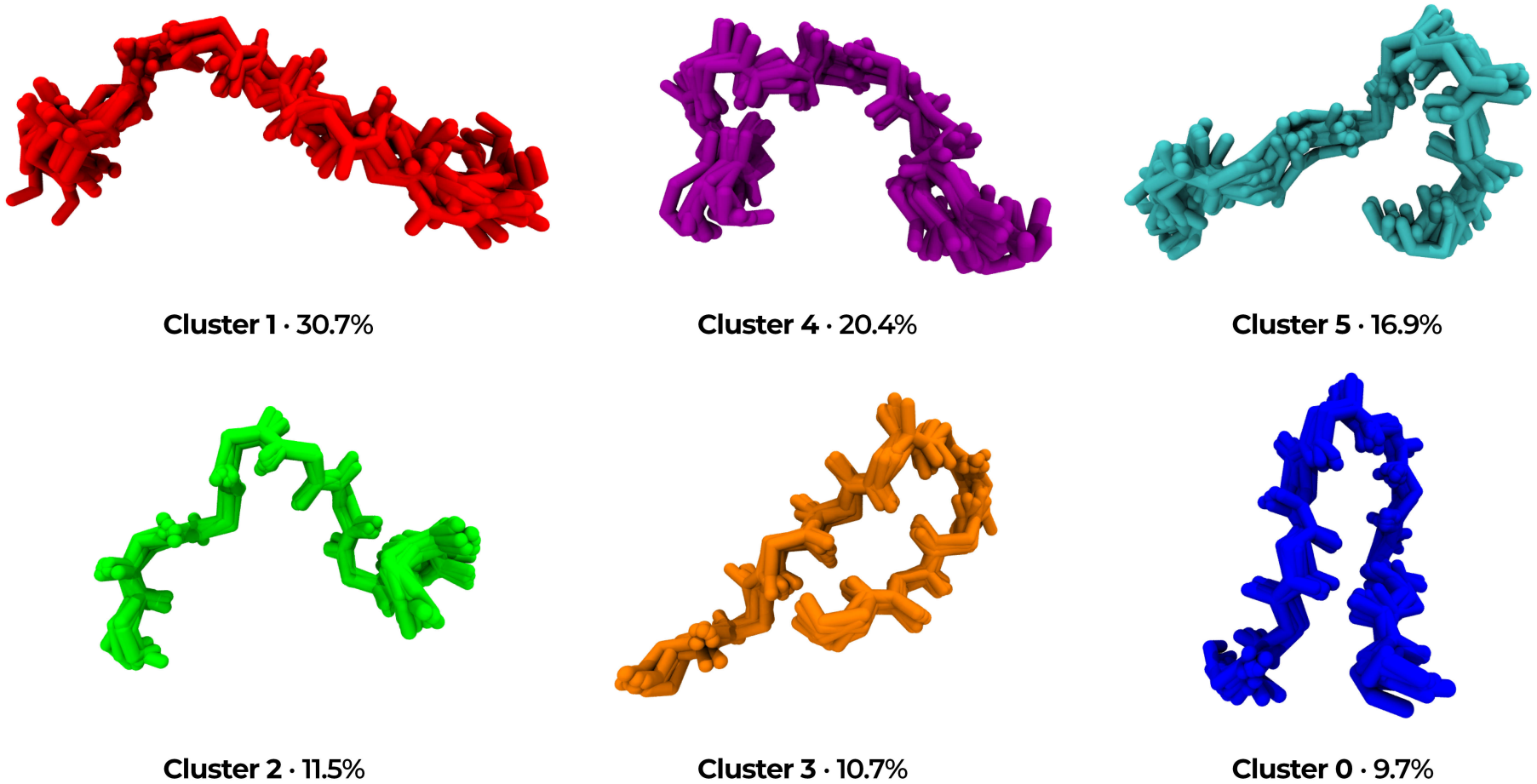
Representative structures of the six tau peptide clusters from the *k* = 6 *k*-means NANI partition (`comp_sim` seeding), rendered in VMD and colored as in the VMDance plots. Each panel shows the cluster medoid overlaid with the nine frames closest to it.

## 5 Conclusions

VMDance brings the MDANCE clustering methods into VMD and keeps cluster assignments, quality metrics, and representative structures tied to the loaded trajectory. Because these methods scale linearly with the number of frames and support deterministic initialization, partitions can be reproduced exactly and recomputed and compared interactively without leaving the viewer. Conformational states are thus identified, evaluated, and inspected where the structures themselves are viewed.

## Data and Software Availability

VMDance is freely available at https://github.com/diegoenry/vmd-mdance-plugin. Documentation, including installation instructions, a tutorial, and a description of the scripting command, is available at https://diegoenry.github.io/vmd-mdance-plugin/. The C++ backend, CPP-MDANCE, is available at https://github.com/mqcomplab/CPP-MDANCE, and the original Python implementation, MDANCE, at https://github.com/mqcomplab/MDANCE. VMDance requires a VMD installation with Tcl/Tk 8.6 and a compiled CPP-MDANCE backend.

## Notes

The authors declare no competing financial interest.

## Acknowledgment

R.A.M.Q., J.B.W.S., K.A.Z., V.J., and J.C.Q. acknowledge support from the National Institute of General Medical Sciences of the National Institutes of Health under award number R35GM150620. D.E.B.G. and R.C.B. acknowledge support from the National Science Foundation under award number MCB-2143787. VMD development is supported by the National Institute of General Medical Sciences of the National Institutes of Health under award number R24-GM145965.

Anthropic’s Claude Opus 5 was used to assist in writing the plugin’s graphical interface and test suite, together with the command-line interface and C API of the CPP-MDANCE backend. Claude was also used to edit the language of the manuscript and Supporting Information. All code and text were reviewed, tested, and validated by the authors, who take full responsibility for the content of this work.

## Supporting Information

Test system and atom selection (Section S1); full elbow scan of *k*-means NANI for *k* = 3–30 (Table S1) and Calinski–Harabasz and Davies–Bouldin indices versus *k* (Figure S1); per-cluster results of the *k* = 6 partition (Table S2); comparison with the Python implementation of MDANCE (Table S3); diagnostic views of the *k* = 6 partition (Figures S2–S13). (PDF)

## TOC Graphic

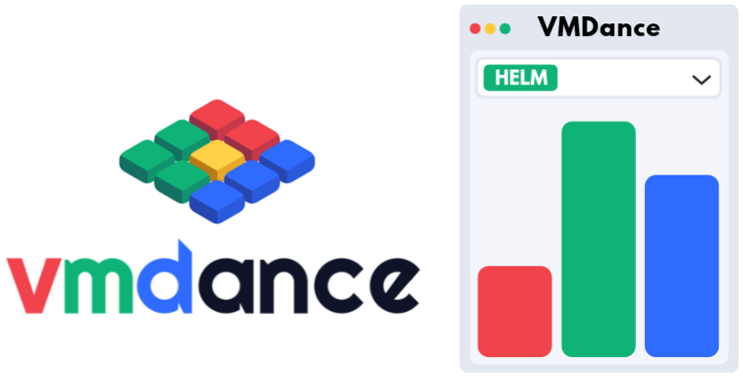

