## Supporting Information for "VMDance: An Efficient and Interactive Molecular Dynamics Clustering Plugin for VMD"

**Contents**

- Section S1. Test system
- Section S2. Selection of the number of clusters
- Section S3. The $k = 6$ partition
- Section S4. Agreement with the Python implementation of MDANCE
- Section S5. Diagnostic views

### S1 Test System

- **System:** capped tau peptide, Ac-GKVQIINKKLDL-$NH_2$ [1–3]
- **Source:** Replica-exchange MD trajectory, 6,001 frames, from the BitQT repository [1;2]
- **Atom selection:** backbone N, C$\alpha$, C, O, and H atoms of residues Lys2–Asp11 (50 atoms; `resid 3 to 12` in the topology file)
- **Alignment and VMD selection string:** `resid 3 to 12 and name N CA C O H`

### S2 Selection of the Number of Clusters

$k$-means NANI (`comp_sim` seeding, 10% seed fraction, MSD metric) was run for $k = 3$–30. The main text uses $k = 3$–10 (Table 2); partitions with $k = 2$ were excluded as trivial splits.

Table S1: Full elbow scan of $k$-means NANI (`comp_sim` seeding, 10% seed fraction) on the tau peptide. CHI, Calinski–Harabasz index (higher is better); DBI, Davies–Bouldin index (lower is better); Avg MSD, average per-cluster mean squared deviation (Å$^2$); Cluster sizes, populations of the smallest and largest clusters. The partition used in the main text is in bold.

| $k$ | CHI | DBI | Avg MSD | Cluster sizes |
|---|---|---|---|---|
| 3 | 1750.28 | 1.8168 | 10.881 | 1326–2338 |
| 4 | 1434.61 | 1.9856 | 10.122 | 1203–1765 |
| 5 | 1298.01 | 1.7952 | 9.026 | 814–1809 |
| **6** | **1142.79** | **1.7809** | **8.349** | **582–1844** |
| 7 | 1051.14 | 1.9452 | 8.107 | 482–1167 |
| 8 | 967.30 | 1.9277 | 7.820 | 415–1097 |
| 9 | 882.01 | 2.0898 | 7.920 | 401–804 |
| 10 | 842.80 | 1.9944 | 7.649 | 457–829 |
| 11 | 780.58 | 1.9499 | 7.142 | 140–826 |
| 12 | 744.84 | 1.9200 | 7.059 | 137–777 |
| 13 | 710.50 | 1.8906 | 6.935 | 137–714 |
| 14 | 684.26 | 1.8692 | 6.928 | 137–718 |
| 15 | 656.41 | 1.8701 | 6.697 | 139–669 |
| 16 | 632.40 | 1.8576 | 6.647 | 138–671 |
| 17 | 608.80 | 1.9178 | 6.626 | 138–528 |
| 18 | 595.60 | 1.9447 | 6.665 | 148–593 |
| 19 | 576.47 | 1.8916 | 6.600 | 150–590 |
| 20 | 563.10 | 1.8400 | 6.487 | 147–586 |
| 21 | 547.60 | 1.8068 | 6.320 | 133–583 |
| 22 | 521.89 | 1.8737 | 6.288 | 136–611 |
| 23 | 520.45 | 1.8507 | 6.115 | 138–578 |
| 24 | 512.97 | 1.8273 | 6.006 | 137–587 |
| 25 | 497.53 | 1.8302 | 5.975 | 126–582 |
| 26 | 487.43 | 1.8316 | 5.940 | 133–579 |
| 27 | 468.78 | 1.8698 | 5.888 | 57–576 |
| 28 | 462.96 | 1.8243 | 5.810 | 56–577 |
| 29 | 447.54 | 1.8837 | 5.847 | 48–585 |
| 30 | 436.84 | 1.9141 | 5.831 | 47–586 |

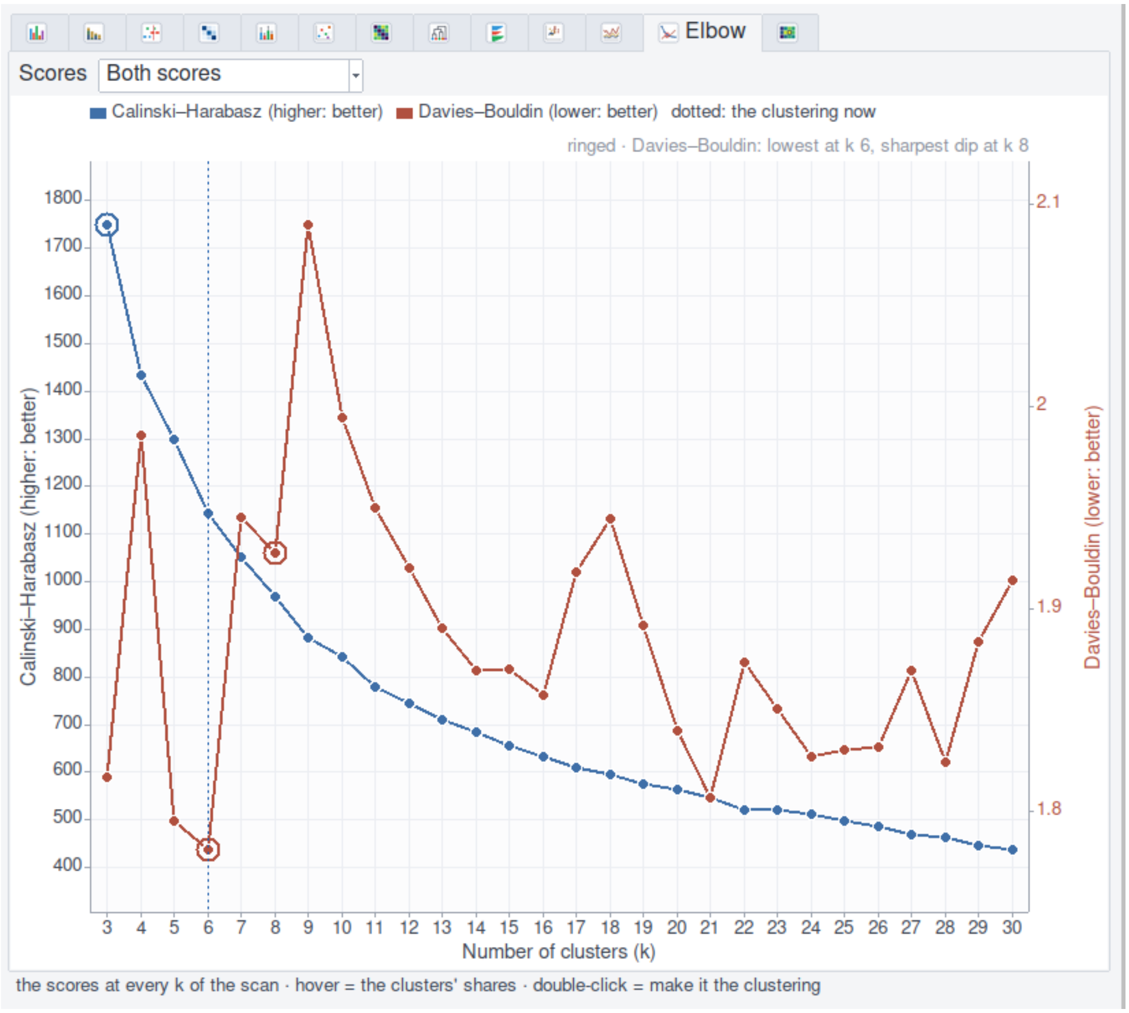


Figure S1: Calinski–Harabasz index (blue, left axis) and Davies–Bouldin index (red, right axis) versus the number of clusters for $k$-means NANI (`comp_sim`) on the tau peptide over $k$ = 3–30, exported from the VMDance Elbow view. Rings mark the highest CHI ($k = 3$), the lowest DBI ($k = 6$), and the sharpest DBI dip, i.e., the largest second derivative ($k = 8$); the dotted line marks the partition used in the main text ($k = 6$).

### S3 The $k = 6$ $k$-means NANI Partition

Table S2: Per-cluster results of the $k = 6$ $k$-means NANI partition (`comp_sim` seeding, 10% seed fraction) as reported by VMDance (cluster numbering as in Figures 2 and 3 of the main text). MSD, per-cluster mean squared deviation; To rep., mean squared deviation to the representative frame; Rep., representative (medoid) frame.

| Cluster | Frames | % | MSD (Å$^2$) | To rep. (Å$^2$) | Rep. |
|---:|---:|---:|---:|---:|---:|
| 1 | 1844 | 30.7 | 10.35 | 8.017 | 5982 |
| 4 | 1225 | 20.4 | 9.677 | 7.273 | 108 |
| 5 | 1012 | 16.9 | 10.83 | 8.536 | 5469 |
| 2 | 693 | 11.5 | 7.758 | 4.738 | 530 |
| 3 | 645 | 10.7 | 4.154 | 2.770 | 2633 |
| 0 | 582 | 9.7 | 7.328 | 5.525 | 2629 |

### S4 Agreement with the Python Implementation of MDANCE

Table S3: Comparison of the $k = 6$ $k$-means NANI partition (`comp_sim` seeding, 10% seed fraction) obtained with the CPP-MDANCE backend and with the Python implementation of MDANCE on identical input.

| Quantity | CPP-MDANCE | MDANCE (Python) |
|---|---:|---:|
| CHI | 1142.79 | 1142.79 |
| DBI | 1.7809 | 1.7809 |
| Avg MSD (Å$^2$) | 8.349 | 8.349 |
| Cluster sizes | 1844, 1225, 1012, 693, 645, 582 | 1844, 1225, 1012, 693, 645, 582 |
| Per-cluster MSD (Å$^2$) | 10.35, 9.68, 10.83, 7.76, 4.15, 7.33 | 10.35, 9.68, 10.83, 7.76, 4.15, 7.33 |

For $k = 3$–15, the CHI, DBI, and average MSD values obtained with the two implementations are also identical to the precision reported in Table S1.

### S5 Diagnostic Views

The figures below show the diagnostic views of VMDance for the $k = 6$ $k$-means NANI partition (`comp_sim` seeding, 10% seed fraction) of the tau peptide, captured from the plugin's plot pane. Cluster numbering and colors are as in Figures 2 and 3 of the main text. The Elbow view is shown in Figure S1.

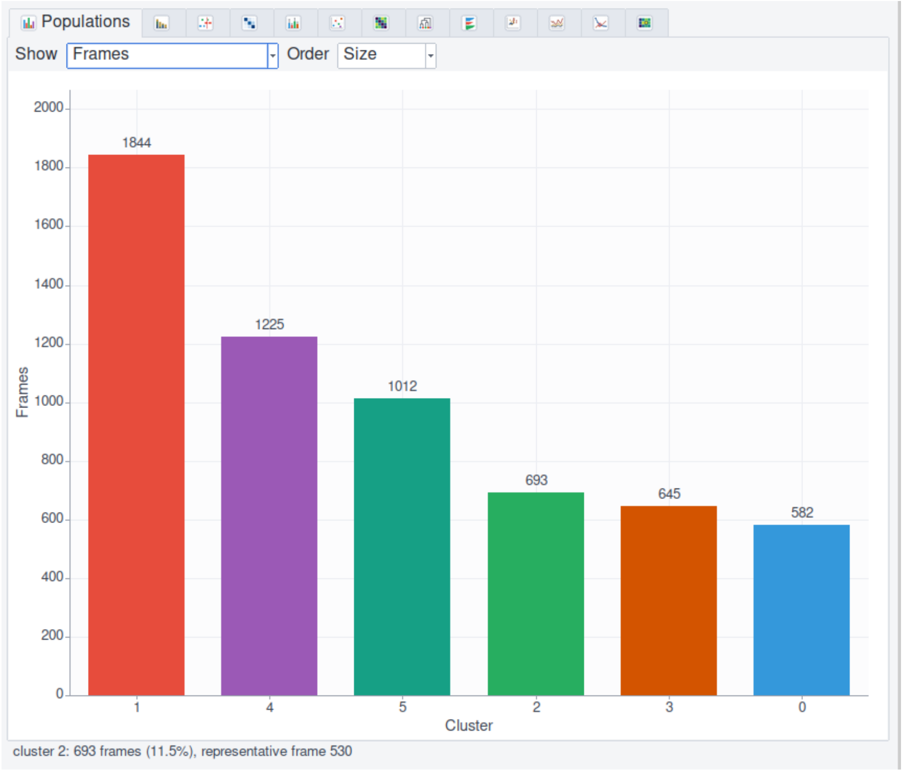


Figure S2: Populations view. Number of frames in each cluster, ordered by size.

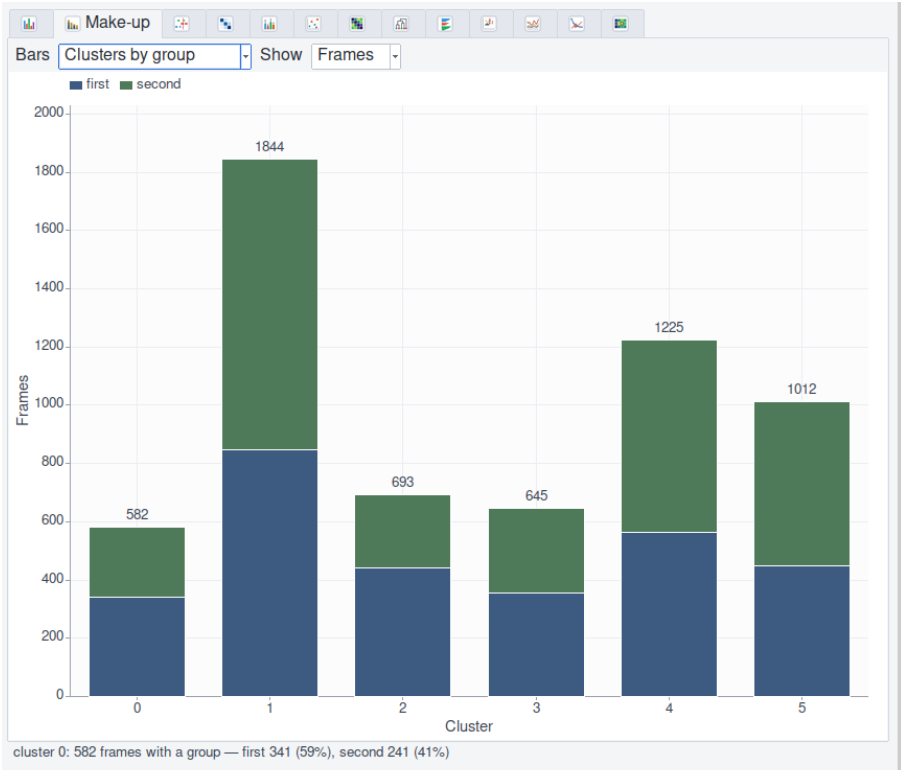


Figure S3: Make-up view. Number of frames in each cluster split by user-defined frame groups, here the first (frames 0–2999) and second (frames 3000–6000) halves of the trajectory.

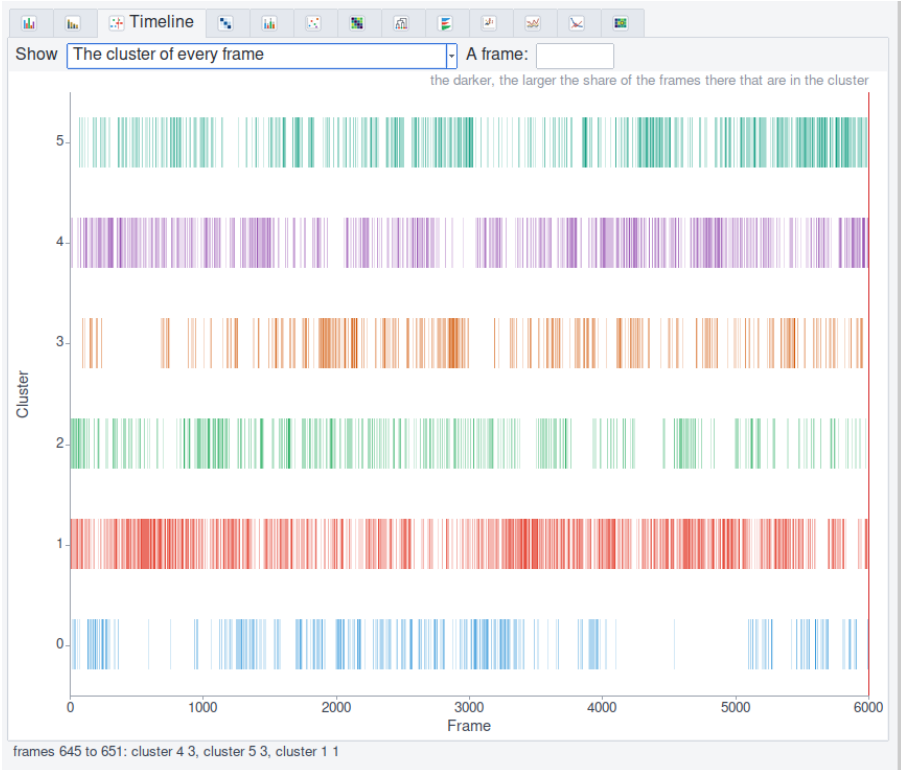


Figure S4: Timeline view. Cluster assignment of every frame along the trajectory.

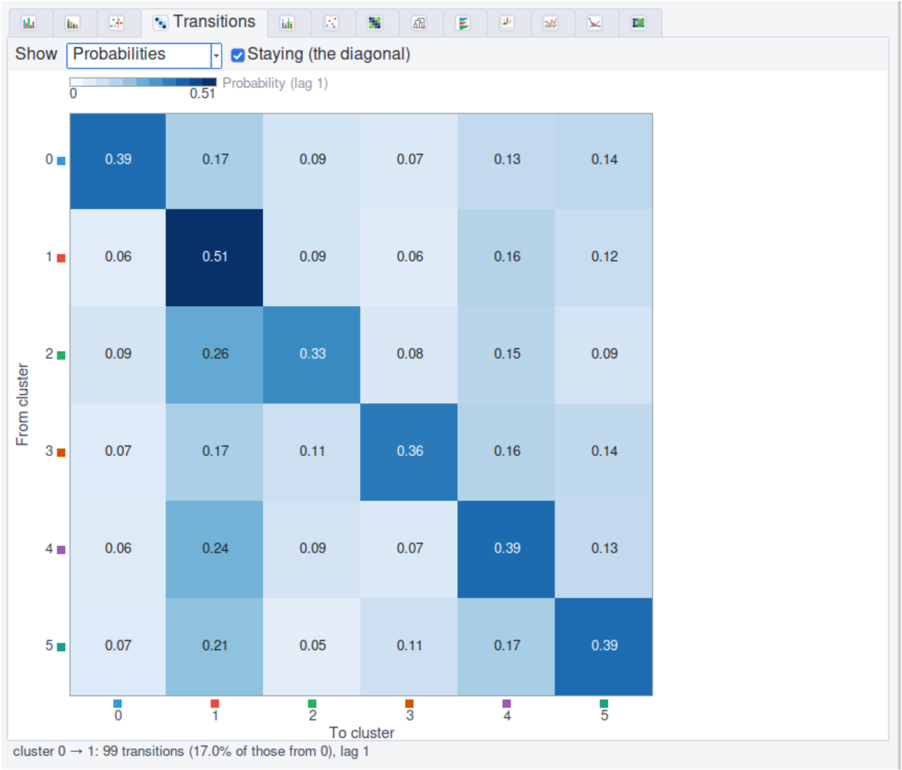


Figure S5: Transitions view. Probabilities of transitions between clusters at a lag of one frame; the diagonal gives the probability of remaining in the same cluster.

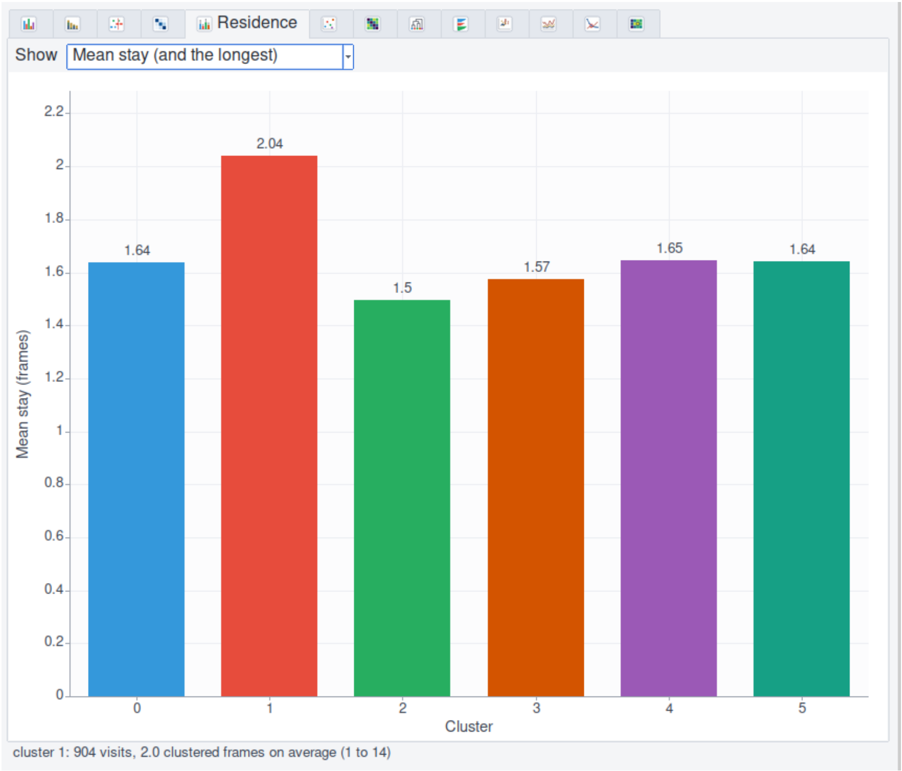


Figure S6: Residence view. Mean number of consecutive frames spent in each cluster per visit.

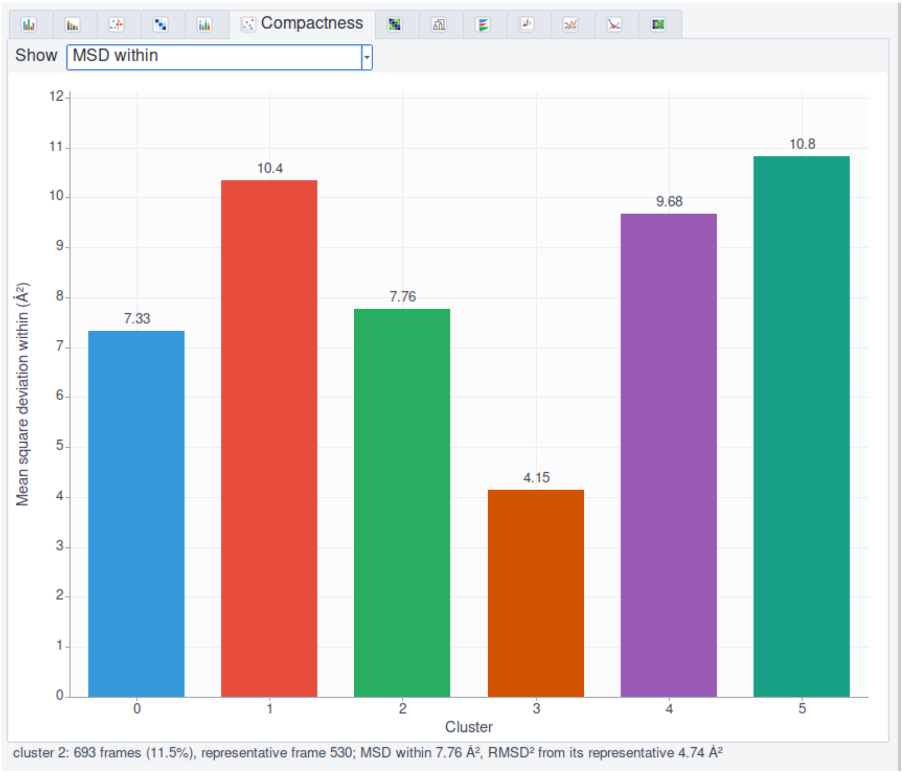


Figure S7: Compactness view. Mean squared deviation within each cluster (Å$^2$).

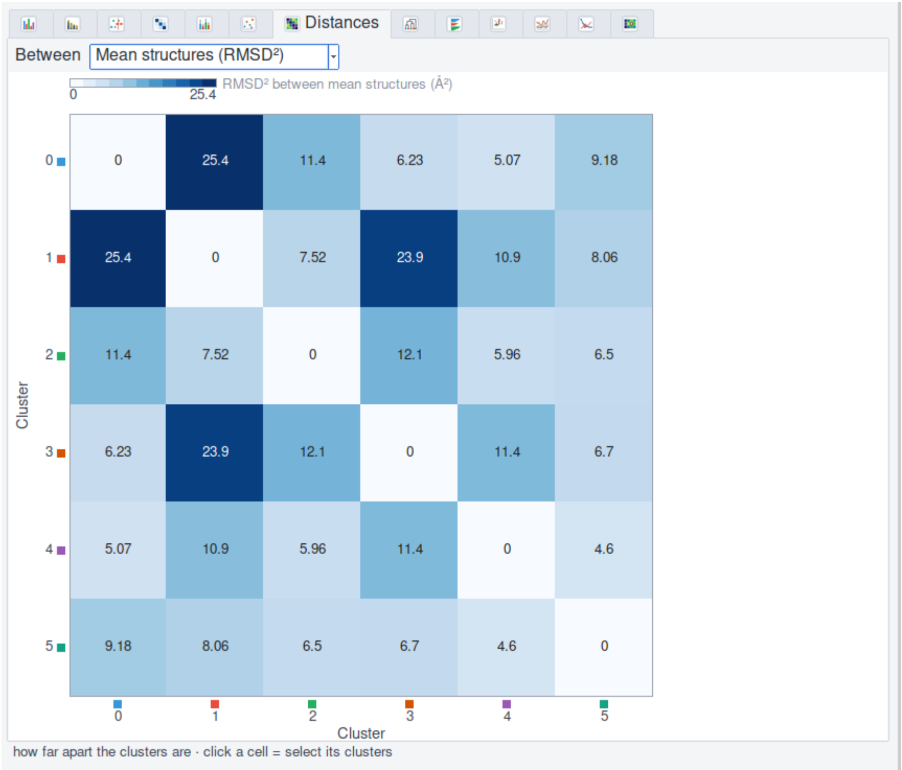


| | 0 | 1 | 2 | 3 | 4 | 5 |
|---|---|---|---|---|---|---|
| 0 | 0 | 25.4 | 11.4 | 6.23 | 5.07 | 9.18 |
| 1 | 25.4 | 0 | 7.52 | 23.9 | 10.9 | 8.06 |
| 2 | 11.4 | 7.52 | 0 | 12.1 | 5.96 | 6.5 |
| 3 | 6.23 | 23.9 | 12.1 | 0 | 11.4 | 6.7 |
| 4 | 5.07 | 10.9 | 5.96 | 11.4 | 0 | 4.6 |
| 5 | 9.18 | 8.06 | 6.5 | 6.7 | 4.6 | 0 |

Figure S8: Distances view. Squared RMSD between the mean structures of the clusters (Å$^2$).

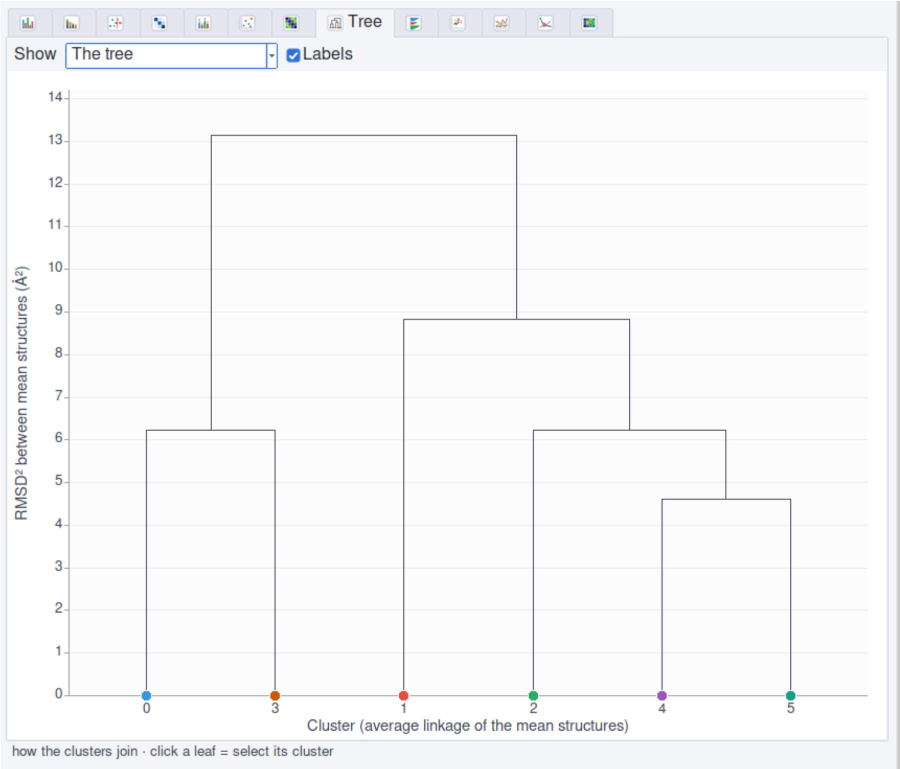


Figure S9: Tree view. Average-linkage dendrogram of the cluster mean structures, with heights given by the squared RMSD between mean structures (Å$^2$).

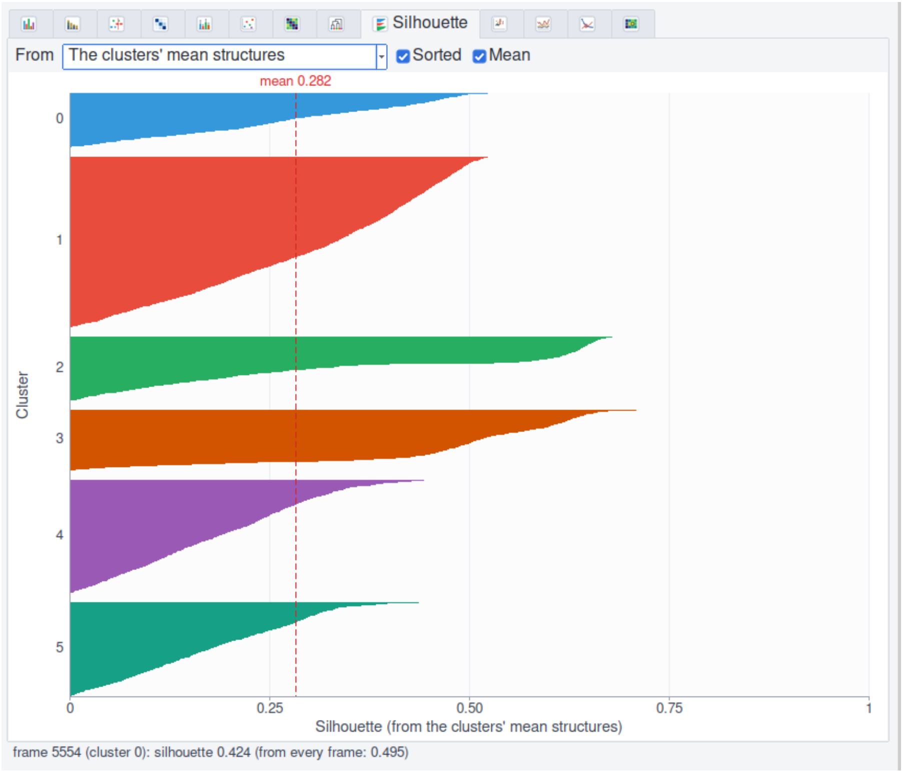


Figure S10: Silhouette view. Simplified silhouette value of every frame computed from the clusters' mean structures, sorted within each cluster; the dashed line marks the mean (0.282).

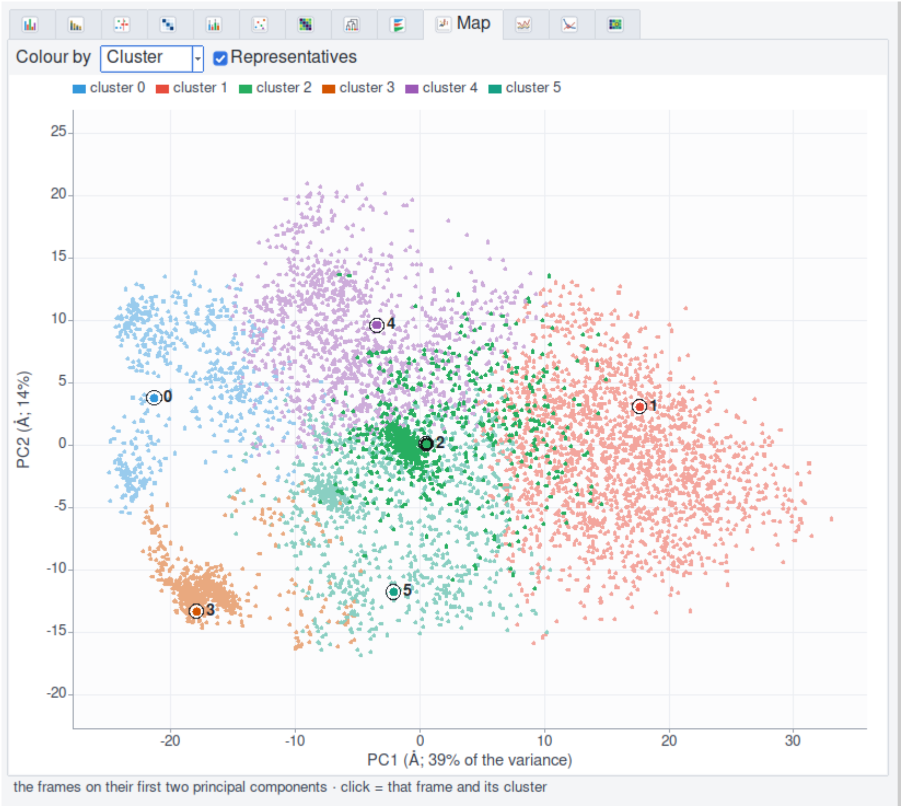


Figure S11: Map view. Projection of every frame onto the first two principal components of the coordinates (PC1, 39% and PC2, 14% of the variance; axes computed from 500 frames), colored by cluster. Circles mark the representative frame of each cluster.

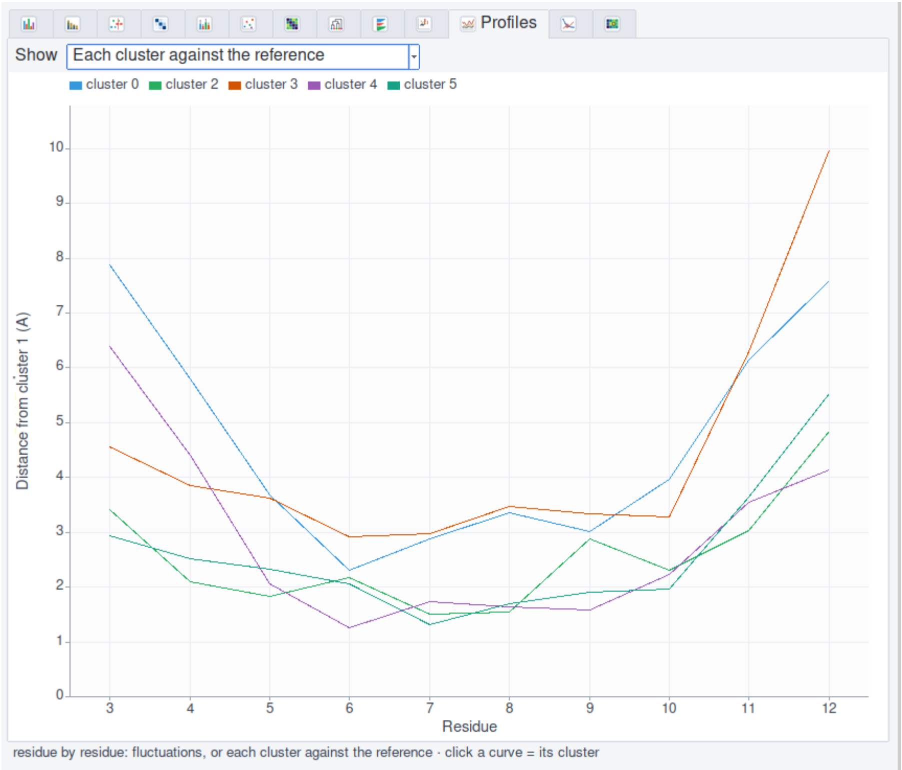


Figure S12: Profiles view. Per-residue distance of the mean structure of each cluster from that of the most populated cluster (cluster 1), for residues 3–12.

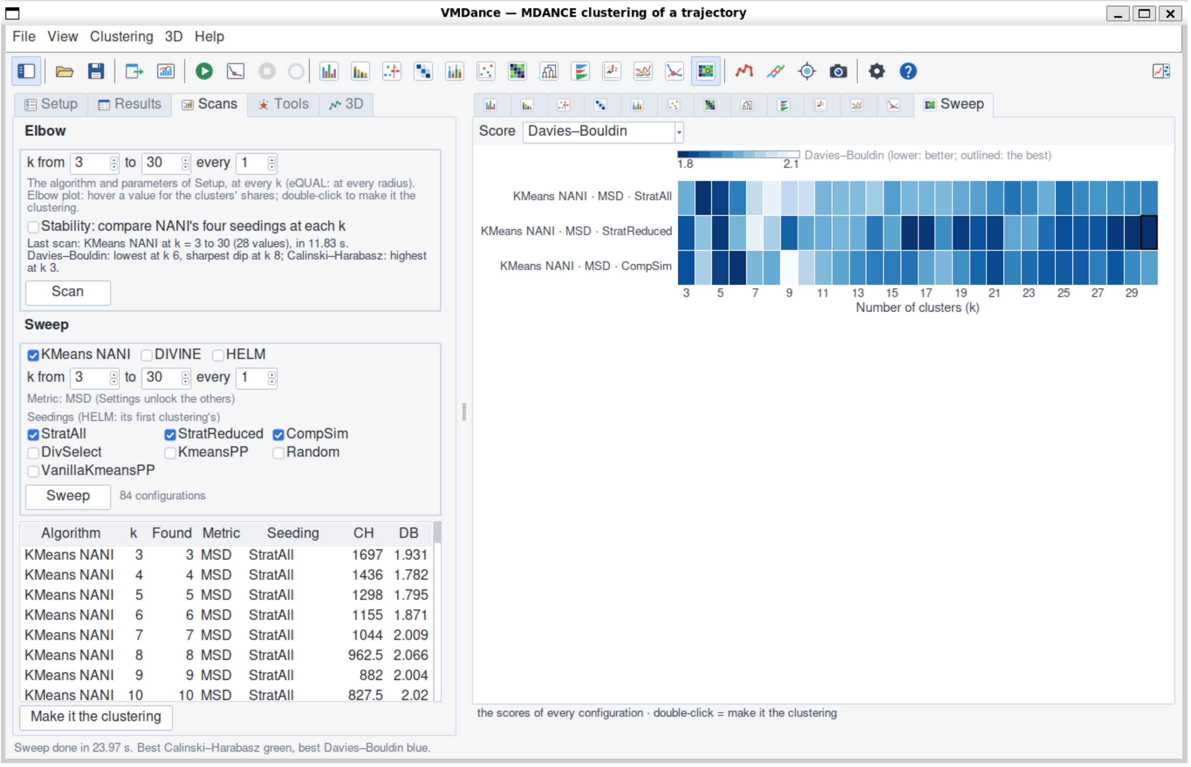


Figure S13: Sweep view. Davies–Bouldin index of $k$-means NANI with `strat_all`, `strat_reduced`, and `comp_sim` seeding over $k = 3$–30, shown as a heat map (right) and as a table of every configuration (left). The full interface is shown.
